# Endothelial Rbpj is essential for the education of tumour-associated macrophages

**DOI:** 10.1101/2021.12.20.473423

**Authors:** Ronja Mülfarth, Elisenda Alsina-Sanchis, Iris Moll, Sarah Böhn, Lena Wiedmann, Lorea Jordana, Tara Ziegelbauer, Jacqueline Taylor, Francesca De Angelis Rigotti, Adrian Stögbauer, Benedetto Daniele Giaimo, Adelheid Cerwenka, Tilman Borggrefe, Andreas Fischer, Juan Rodriguez-Vita

## Abstract

Epithelial ovarian cancer (EOC) is one of the most lethal gynaecological cancers worldwide. EOC cells educate tumour-associated macrophages (TAMs) through CD44-mediated cholesterol depletion to generate an immunosuppressive tumour microenvironment (TME). In addition, tumour cells frequently activate Notch1 receptors on endothelial cells (ECs) to facilitate metastasis. However, little is known whether the endothelium would also influence the education of recruited monocytes. Here, we report that canonical Notch signalling through RBPJ in ECs is an important player in the education of TAMs and EOC progression. Deletion of *Rbpj* in the endothelium of adult mice reduced infiltration of monocyte-derived macrophages into the TME of EOC and prevented the acquisition of a typical TAM gene signature. This was associated with stronger cytotoxic activity of T cells and decreased tumour burden. Mechanistically, we identified CXCL2 as a novel Notch/RBPJ target gene. This angiocrine factor regulates the expression of CD44 on monocytes and subsequent cholesterol depletion of TAMs. Bioinformatic analysis of ovarian cancer patient data showed that increased CXCL2 expression is accompanied by higher expression of CD44 and TAM education. As such, EOC cells employ the tumour endothelium to secrete CXCL2 in order to facilitate an immunosuppressive microenvironment.

## Introduction

High grade serous ovarian cancer is the deadliest type of all gynaecological cancers^1^. The high mortality rate is due to the fact that most women have already developed peritoneal metastasis at the time of diagnosis. Epithelial ovarian cancer (EOC) cells can directly infiltrate the peritoneal cavity to seed metastases, a process called transcoelomic metastasis^2^. Metastatic EOC cells initially reside in the omentum, where they undergo certain adaptations allowing them to spread throughout the whole peritoneal cavity. Importantly, this peritoneal microenvironment is so immunosuppressive that even the infiltration of effector T cell does not *per se* correlate with better prognosis^3^.

Peritoneal spread of tumour cells is accompanied by monocyte-derived macrophage (MN-derived macrophages) recruitment, which eventually become the most abundant myeloid cell type^4^ and are a major contributor to the immunosuppressive TME in EOC^3^. Upon recruitment from blood to the tumour, monocytes differentiate into macrophages which are further educated by the TME. Eventually, tumour-associated macrophages (TAMs) strongly promote the progression of metastatic ovarian cancer^5^. However, little is known about the contribution of other stromal cells to the peritoneal spread of EOC cells and to macrophage education.

Monocytes must cross the vascular endothelial barrier before infiltrating peritoneal organs or the peritoneal fluid. Endothelial cells (ECs) not only form tubes for the transport of blood, but also produce soluble factors controlling the differentiation and function of adjacent cells. These angiocrine functions are highly context and organ specific^6,7^. In cancer, ECs provide angiocrine factors which influence tumour progression^8^. Therefore, we hypothesized that monocytes are influenced by ECs, for instance during transmigration, while infiltrating into peritoneal tissues.

Notably, tumours can alter the transcriptome of ECs^9,10^ and this may also influence transmigrating monocytes. For example, endothelial Notch signalling activity is frequently higher in tumours like EOC and in the metastatic niche compared to ECs from non-tumourous tissue^11^. Notch signalling is a highly conserved cell-to-cell communication system. Ligand binding induces cleavage of the transmembrane Notch receptors releasing the intracellular domain (ICD) which enters the nucleus to alter gene transcription. This canonical signalling pathway relies on the DNA-binding protein RBPJ, which turns into a transcriptional activator upon binding of a Notch receptor ICD^12^. Sustained endothelial Notch1 signalling is associated with increased myeloid cell infiltration and metastasis^11^. The Notch pathway in ECs is a major regulator of angiogenesis, metabolism, angiocrine functions and tumour cell transmigration^11,13-18^. Although EOC cells do not necessarily have to cross the endothelial barrier to spread throughout the peritoneum we hypothesized that endothelial Notch signalling could still influence EOC progression via transmigrating myeloid cells.

Here, we provide insights into the essential role of RBPJ in ECs for the recruitment of monocytes to the microenvironment of metastatic EOC and their proper education into pro-tumoural TAMs.

## Results

### Deletion of *Rbpj* in endothelial cells reduces EOC progression

Metastatic EOC cells seed initially in the omentum and later spread within the peritoneum. During these steps, tumour cells undergo transcriptional changes that allow them to further grow and colonise (**Fig. 1a**). The latter is strongly influenced by MN-derived macrophages^19^. To determine the contribution of canonical Notch signalling in ECs to myeloid cell infiltration and EOC progression, we used the tamoxifen-inducible VE-Cadherin (Cdh5) Cre^ERT2^ strain to delete *Rbpj* specifically in ECs of adult mice (*Rbpj*^iΔEC^). This mouse model is well established and leads to robust gene recombination in ECs of several organs^14,15,17,18^ (**Fig. 1b**). Under physiological conditions, there were no differences in blood vessel density in the larger omentum upon *Rbpj* deletion in ECs compared to controls (**Suppl. Fig. 1a,b**). However, upon intraperitoneal injection of ID8 cells mimicking metastatic EOC, the omenta showed evidence of tumour nodule growth (**Fig. 1c**) and had significantly higher vessel density in *Rbpj*^iΔEC^ mice compared to control animals (**Fig. 1d**). The density of endothelial coverage with α-smooth muscle actin-positive mural cells was, however, unchanged (**Suppl. Fig. 1c**). Despite increased vessel density, tumour burden in omenta of *Rbpj*^iΔEC^ mice was significantly lower than that in their littermate controls (**Fig. 1e**). Moreover, peritoneal spread of tumour cells was significantly reduced as well in *Rbpj*^iΔEC^ mice (**Fig. 1f**) concluding that deletion of *Rbpj* in ECs reduces EOC progression and metastasis.

**Figure 1.**
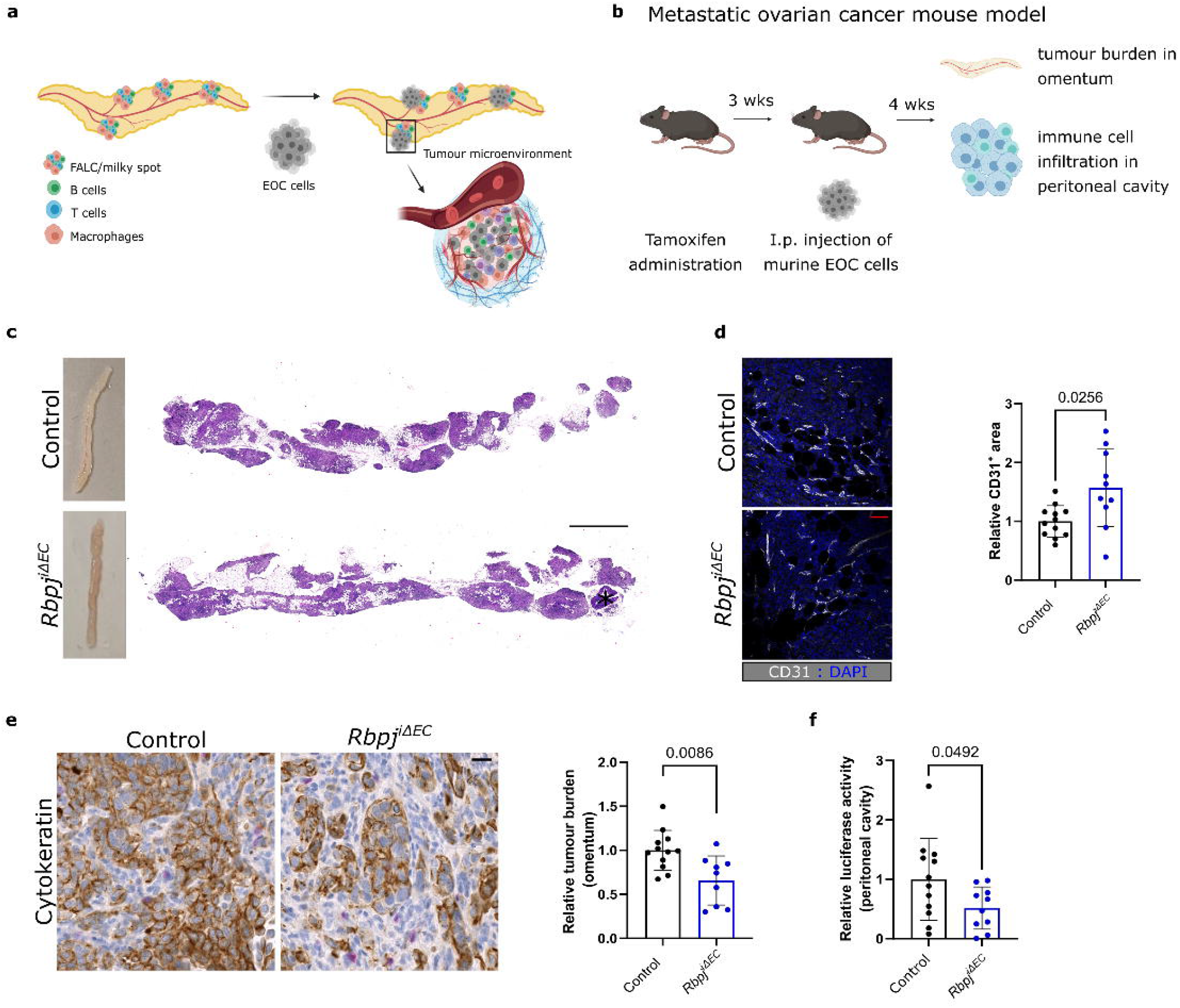

### Deletion of endothelial *Rbpj* decreases monocyte-derived macrophage recruitment

Next, we analysed the immune cell compartment of our EOC model since it is a major contributor to tumour progression. High Notch signalling activity in ECs induces VCAM1 expression, which promotes leukocyte extravasation^11,20^. However, VCAM1 expression was unchanged between *Rbpj*^iΔEC^ and their controls littermates (**Suppl. Fig. 2a**). Nevertheless, whole-mount staining of omenta revealed that there was a reduction in immune cells inside tumour nodules of *Rbpj*^iΔEC^ mice compared to controls (**Fig. 2a** and **Suppl. Fig. 2b**). Interestingly, tumours in control mice contained more cells with a large vacuolated cytoplasm, reminiscent of macrophages. Therefore, we analysed this cell population in greater detail. Tumour-bearing omenta from *Rbpj*^iΔEC^ mice contained less CD11b^+^ cells within tumour nodules, although this reduction was not statistically significant (**Fig. 2b**). In order to better determine the macrophage subpopulations responsible for the observed decrease in CD11b^+^ cells; we studied the myeloid and macrophage populations in the peritoneal cavity to understand whether RBPJ in ECs could play a role in their composition. After four weeks of tumour growth, there were no significant differences in the total amount of myeloid cells or macrophages (**Fig. 2c,d**). However, we found decreased numbers of small peritoneal macrophages (SPMs), which are MN-derived macrophages characterized as MHC-II^hi^/F4/80^low^ (**Fig. 2e**) and CCR2^+^/Tim4^-^ (**Fig. 2f**). Notably, in naïve (tumour-free) conditions, endothelial *Rbpj* deletion had no effect on peritoneal macrophage populations (**Suppl. Fig. 2c-f**). These data suggest that the tumour-driven recruitment of monocytes into the peritoneum is impaired upon endothelial deletion of *Rbpj*.

**Figure 2.**
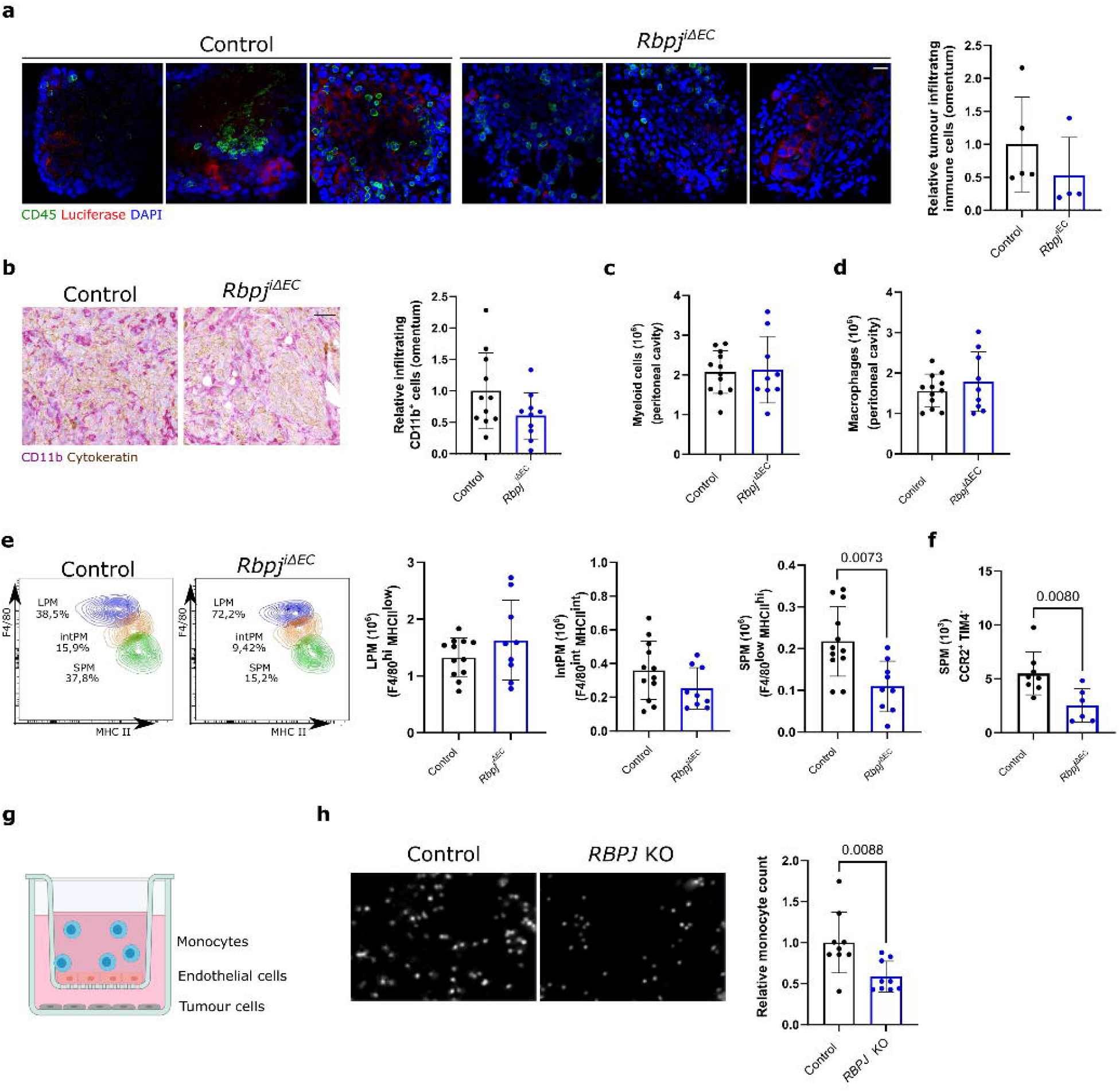

To further dissect endothelial chemotaxis in a more simplified *in vitro* system, we switched to the human system and measured transmigration of human CD14^+^ monocytes through a monolayer of human ECs in a transwell insert. Chemotaxis was stimulated by SK-OV-3 human ovarian cancer cells below the insert (**Fig. 2g**). *RBPJ* deletion in human umbilical vein ECs (HUVECs) resulted in significantly decreased monocyte transmigration rates compared to control HUVECs (**Fig. 2h**). This further suggests that canonical Notch signalling in ECs is important to facilitate monocyte recruitment into the TME.

### Canonical Notch signalling in ECs regulates monocyte recruitment through CXCL2

We have previously analysed the transcriptional induction of chemokines in HUVECs upon Notch1 ICD (N1ICD) overexpression^11^. Based on this, we transduced *RBPJ*-deficient HUVEC and control cells with N1ICD to determine which of those transcriptional changes require *RBPJ*. The chemokines that were induced by N1ICD through RBPJ were *CCL1, CCL21, CXCL12*, and *CXCL2* (**Fig. 3a**) and we excluded those that were not induced through RBPJ (**Suppl. Fig. 3a-b**). Next, we measured the mRNA expression levels of these chemokines in peritoneal adipose tissue obtained from *Rbpj*^iΔEC^ mice. This revealed that there was lower *Cxcl2* expression in peritoneal adipose tissue from *Rbpj*^iΔEC^ mice compared to littermate controls, whereas the other chemokines analysed were not changed (**Fig. 3b**). Analysis of the same cytokines in endothelial-specific N1ICD mice^11,15^, exposed that higher endothelial Notch1 signalling activity led to higher *Cxcl2* expression in peritoneal adipose tissue (**Fig. 3c**).

**Figure 3.**
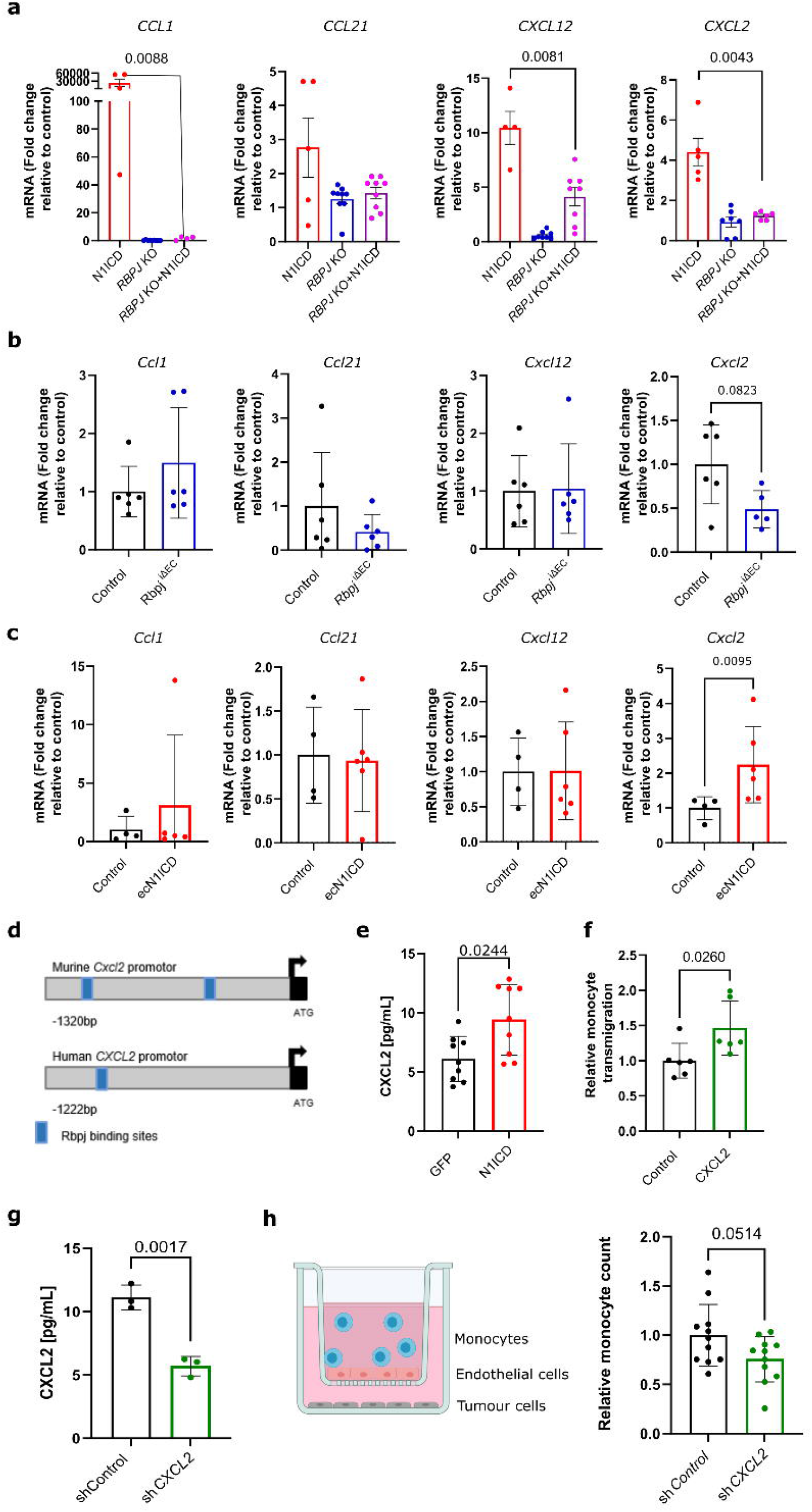

*In silico* analysis showed that the mouse and human *Cxcl2* gene contains RBPJ-binding sites in the promoter region (**Fig. 3d**). Consistently, cultured human ECs secreted higher CXCL2 protein levels after being transduced with N1ICD expressing adenovirus compared to GFP-transduced control cells (**Fig. 3e**).

These data revealed that canonical Notch signalling induces CXCL2 expression in ECs. CXCL2 is known to attract granulocytes but also to a lesser extend monocytes^21^. As such, it poses as an interesting endothelial Notch target, which could mediate the observed effects on monocyte recruitment into the TME. To evaluate whether CXCL2 was capable of attracting monocytes, we performed transwell chemotaxis experiments and observed that recombinant CXCL2 induced monocyte chemotaxis (**Fig. 3f**). Next, we silenced *CXCL2* expression in ECs using shRNA, which led to an about 50% reduction of CXCL2 protein expression (**Fig. 3g**). Compared to non-silencing control, this reduction of CXCL2 levels was already capable of reducing the numbers of monocytes transmigrating through ECs towards SK-OV-3 cells (**Fig. 3h**). In summary, the data implicate that the endothelial RBPJ/CXCL2 axis contributes to monocyte recruitment into the peritoneum during transcoelomic metastasis.

### *Rbpj* in ECs is necessary for tumour-mediated TAM education

Recruited monocytes differentiate into macrophages which are further educated by tumour cells to become tumour-promoting macrophages. Macrophage education in EOC is facilitated by hypersensitivity towards IL4, which is induced by tumour cell-mediated cholesterol depletion^4^. We sought to understand whether lack of *Rbpj* in ECs could influence macrophage phenotypes. To assess this, we isolated newly recruited MN-derived macrophages (CD11b^+^/F4/80^+^/CCR2^+^) (**Fig. 4a**) and obtained their transcriptomic profile by microarray analysis. Gene set enrichment analysis (GSEA) comparing MN-derived macrophages from *Rbpj*^iΔEC^ and control tumour-bearing mice revealed that *Rbpj* in ECs is necessary to acquire the typical phenotype of TAM in this model of metastatic EOC (**Fig. 4b**). As such, tumour cells could not fully educate MN-derived macrophages in mice lacking endothelial *Rbpj*.

**Figure 4.**
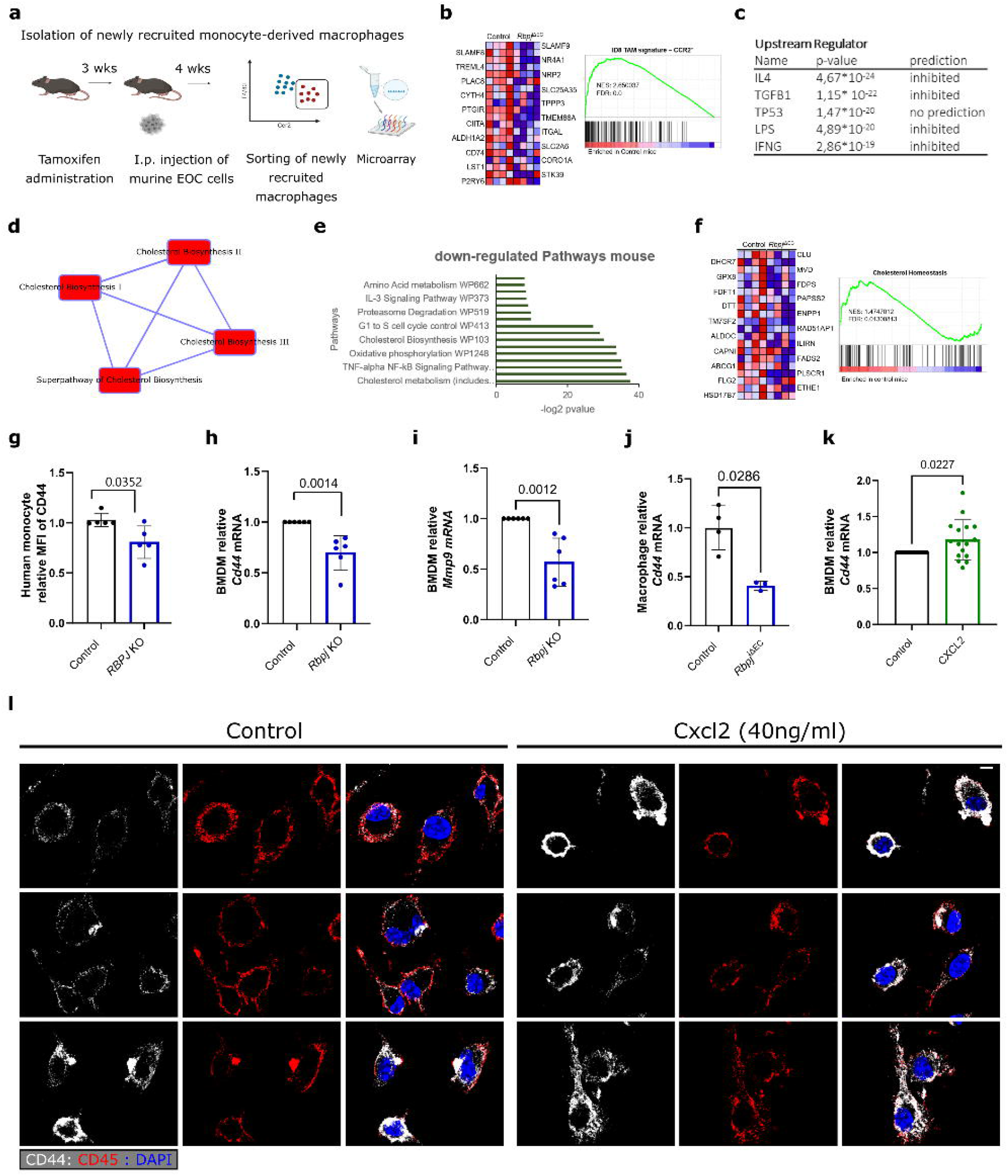

Transcriptomic profiling via Ingenuity Pathway Analysis (IPA) determined that IL4 was the most downregulated signalling pathway in MN-derived macrophages obtained from *Rbpj*^iΔEC^ tumour bearing mice (**Fig. 4c**). IPA and pathway analysis showed that genes important for cholesterol synthesis were downregulated in MN-derived macrophages from *Rbpj*^iΔEC^ mice (**Fig. 4d,e**). We then employed a cholesterol homeostasis gene set from TAMs obtained 21 days after tumour injection^4^, representing EOC-induced cholesterol metabolism in TAMs. GSEA showed that this gene set was significantly enriched in newly recruited macrophages coming from control tumour-bearing mice (**Fig. 4f**), indicating that *Rbpj* in ECs is necessary for cholesterol depletion in TAMs.

Cholesterol depletion is mediated by tumour cell-secreted high molecular weight hyaluronan (HMW-HA). The interaction with HA receptors, such as CD44, in macrophages leads to cholesterol efflux through ABC transporters^4^. In order to understand whether *Rbpj* in ECs could be important for CD44 expression in monocytes, we co-cultured human monocytes with HUVECs lacking *RBPJ* or with respective controls. Monocytes co-cultured with *RBPJ*-deficient HUVECs, had less CD44 on their membrane than those incubated with control HUVECs (**Fig. 4g**). Next, when incubating bone marrow-derived macrophages (BMDMs) with conditioned medium (CM) from immortalized mouse cardiac endothelial cells (MCECs) lacking *Rbpj* (CRISPR-Cas9 mediated), macrophages expressed less *Cd44* (**Fig. 4h**) and *Mmp9* (**Fig. 4i**), a known CD44 target gene. This indicates that a secreted angiocrine factor regulated by the transcription factor RBPJ in ECs is necessary for the regulation of CD44 in macrophages. To understand whether this would also happen *in vivo, Rbpj*^iΔEC^ and control mice were injected with thioglycolate to induce MN-derived macrophage recruitment in the absence of tumour cells. We found that MN-derived macrophages in *Rbpj*^iΔEC^ mice expressed significantly less *Cd44* than those in control mice (**Fig. 4j**), confirming that RBPJ in EC is essential for *Cd44* regulation in MN-derived macrophages *in vivo*.

Considering the important role that CXCL2 had in monocyte recruitment in *Rbpj*^iΔEC^tumour bearing mice and that higher level of CXCL2 in serum is associated not only with myeloid cell infiltration into the TME, but also with worse prognosis for EOC patients^22^, we decided to analyse its role in regulating CD44 expression. Indeed, when stimulating BMDMs with CXCL2, *Cd44* expression was increased (**Fig. 4k**). Moreover, since CD44 is a membrane-bound receptor, we analysed by immunofluorescence whether also its cellular localization could be altered. We observed that CXCL2 stimulation of BMDMs increased the presence of CD44 on the plasma membrane, which would consequently increase its accessibility to hyaluronic acid (**Fig. 4l** and **Suppl. Fig. 4**).

### The TAM gene signature is enriched in human ovarian carcinoma samples with high CXCL2 expression

The data presented so far indicate that the angiocrine factor CXCL2, which is under transcriptional control of Notch/RBPJ signalling, preconditions monocytes to be educated by tumour cells in the TME. In order to evaluate the relevance of this, we analysed *CXCL2* and *CD44* mRNA expression levels in publicly available data sets from the Cancer Genome Atlas (TCGA). Remarkably, the stratification of ovarian cancer patients in 25% upper (CXCL2^hi^) and lower (CXCL2^low^) *CXCL2* expression showed that CXCL2^hi^ patients displayed significantly more expression of *CD44* (**Fig. 5a**). These data suggest a positive correlation between *CXCL2* and *CD44* expression in human tumours. Our data indicate that CXCL2-mediated CD44 induction reflects the ability of tumour cells to educate TAMs. Therefore, the correlation in expression between CXCL2 and CD44 should in turn have implications on TAM education. Therefore, we performed GSEA comparing CXCL2^hi^ and CXCL2^low^ patients using the abovementioned TAM signature. We observed that the TAM signature was significantly enriched in CXCL2^hi^ patients (**Fig. 5b**), indicating that the increased CD44 expression in these patients consequently impacts TAM education.

**Figure 5.**
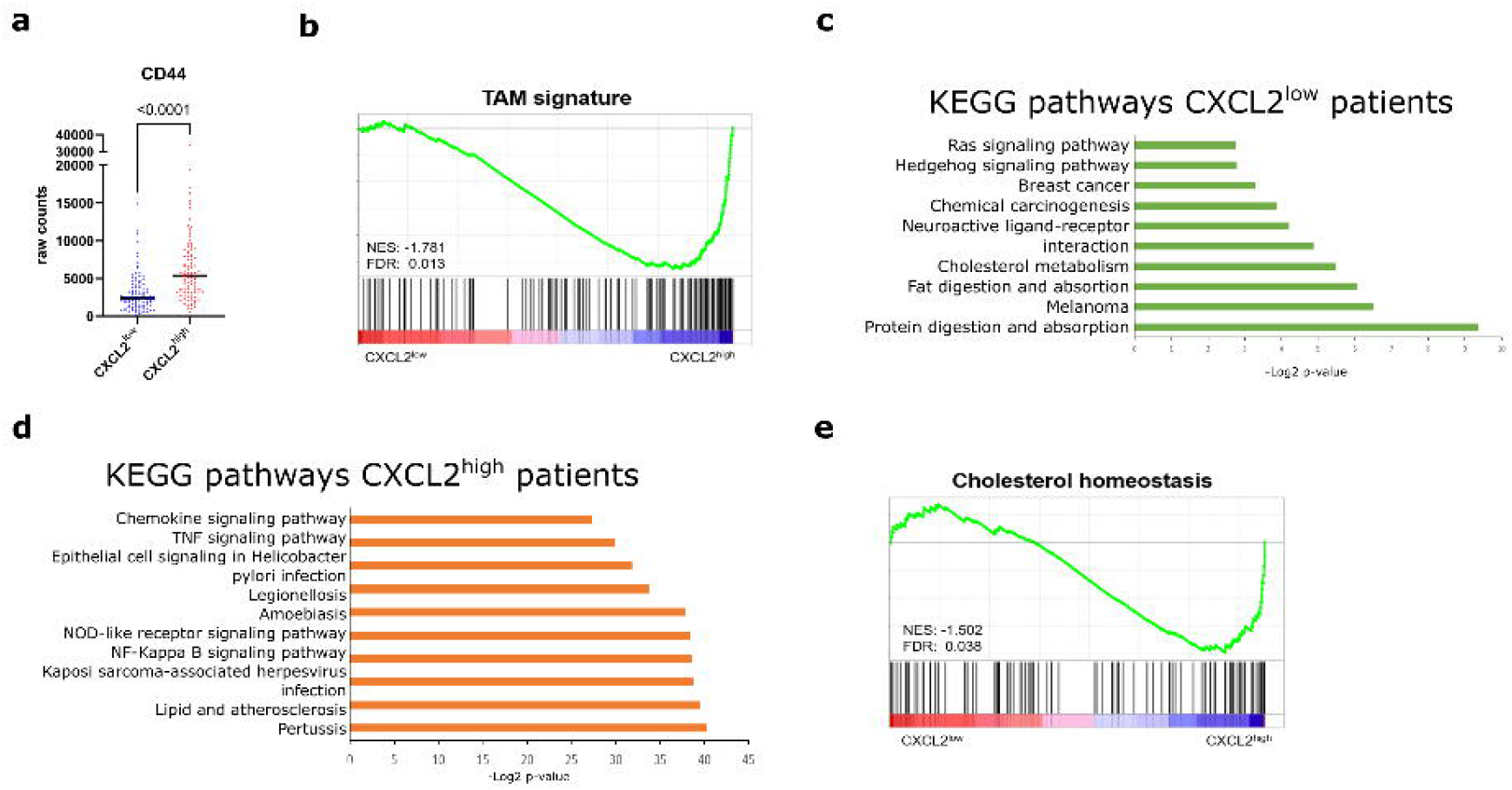

When analysing difference between CXCL2^hi^ and CXCL2^low^ patients, we found an expected upregulation of chemotactic response and myeloid cell recruitment, as shown by gene ontology (GO) term analysis in CXCL2^hi^ patients (**Suppl. Fig. 5**), supporting the role of CXCL2 in controlling the myeloid cell compartment of tumours. Interestingly, we again identified cholesterol metabolism as one of the most downregulated pathways in CXCL2^low^ patients (**Fig. 5c**). Instead, CXCL2^hi^ patients showed induction of pathways such as lipid and atherosclerosis, thus suggesting opposite profiles (**Fig. 5d**). Indeed, GSEA showed that the same cholesterol metabolism gene set previously employed was significantly enriched in CXCL2^hi^ cohort (**Fig. 5e**), indicating a similar cholesterol depletion as the one occurring in peritoneal macrophages of our mouse model.

### TAM education mediated by endothelial *Rbpj* affects T cell cytotoxicity

One of the genes downregulated in MN-derived macrophages from *Rbpj*^iΔEC^ tumour-bearing mice is *Cd74* **(Fig. 4b**). CXCL2 is a ligand for CXCR2^21^, and CD44 is part of a receptor complex that contains CD74 and CXCR2^23,24^. Moreover, CD44 plays an important role in CD74-mediated signal transduction^25^. *In silico* analysis showed that CD74 interacts closely with CD44 and CXCR2 (**Fig. 6a**). Besides, CD74 has been reported to be important in TAMs in brain metastasis^26^, and CD74 expression has been associated with worse prognosis in metastatic ovarian cancer^27^. Furthermore, CD74 was associated with an immunosuppressive phenotype in macrophages^28^. We decided to investigate whether the downregulation of CD74 could have a consequence on TAM behaviour. For that, we analysed publicly available data sets where IL4 responses in CD74-deficient macrophages were analysed^29^. GSEA showed that TAM signature was enriched in wild-type macrophages, indicating that CD74 is not only regulated by IL4, but also necessary for TAM education (**Fig. 6b**). We extracted a signature with the 500 most enriched genes in control compared to CD74-deficient macrophages stimulated with IL4 (CD74-mediated signature), representing a group of genes induced by IL4 through CD74 activation. When comparing newly recruited MN-derived macrophages from our EOC model by GSEA, we found that the CD74-mediated signature was significantly enriched in macrophages isolated from control mice compared to *Rbpj*^iΛEC^. This suggests that newly recruited macrophages from *Rbpj*^iΔEC^ mice cannot induce their immunosuppressive phenotype due to downregulation of CD74-mediated genes (**Fig. 6c**).

**Figure 6.**
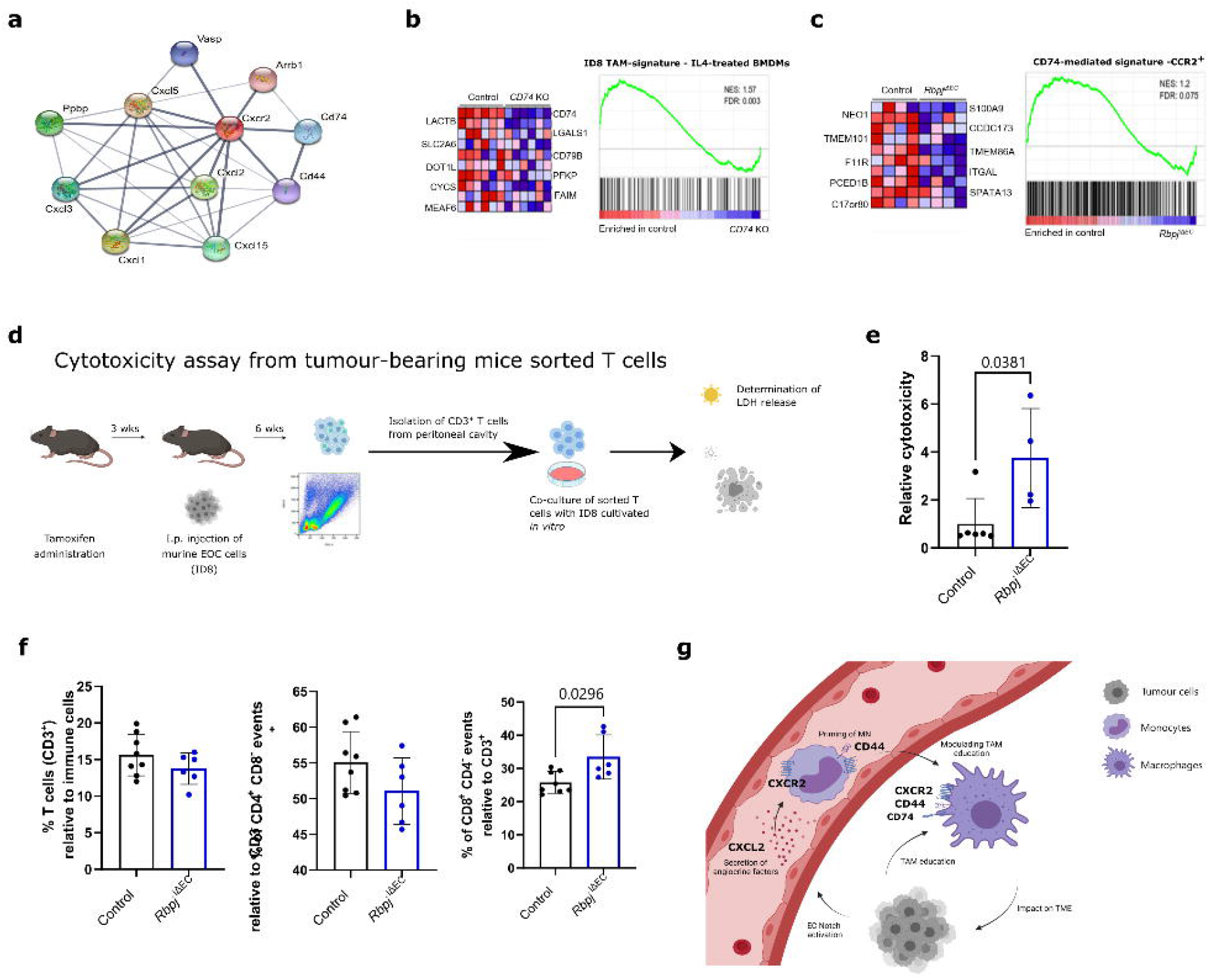

To verify the specific effect on the newly recruited macrophages, we repeated the same analysis on resident macrophages (CD11b^+^/F4/80^+^/CCR2^-^), in which *Cd74* is not differentially expressed between *Rbpj*^iΔEC^ tumour-bearing and control mice, and found no enrichment of this gene set in any group, confirming that *Rbpj* deletion in ECs only affects *Cd74* expression in newly recruited MN-derived macrophages (**Suppl. Fig. 6**). In summary, only macrophages that have crossed the EC barrier as monocytes were affected by the lack of *Rbpj* in the endothelium.

It has been previously reported that TME in metastatic ovarian cancer is highly immunosuppressive and infiltration with immune effector cells has little impact in patients’ outcome^3^. This immunosuppressive microenvironment has been attributed to TAMs^3^. For this reason, we wanted to test whether impaired TAM education could impact on the phenotype of the TME. We isolated T cells from tumour-bearing *Rbpj*^iΔEC^ mice and their littermate controls six weeks after intraperitoneal injection of ID8 cells (**Fig. 6d**). Results demonstrated that T cells derived from *Rbpj*^iΔEC^ mice were more efficient in killing cultured ID8 cells (**Fig. 6e**). This confirms that impairment in TAM education has a direct impact on T cell cytotoxicity. In addition, the analysis of T cell populations in peritoneal cavity from tumour-bearing mice revealed that, although the total frequency of T cells was not changed, the cytotoxic CD8^+^ T cell population was significantly increased in *Rbpj*^iΔEC^ mice compared to their littermate controls (**Fig. 6f**). In summary, our data reveal that *Rbpj* in ECs is necessary for tumour cell-mediated education of MN-derived macrophages into TAMs (**Fig. 6g**).

## Discussion

Collectively, this study provides evidence about a novel angiocrine axis influencing the tumour immune microenvironment. We show how Notch/RBPJ-mediated transcription in ECs, which is frequently hyperactive in tumours^11^, is required for CXCL2-mediated monocyte chemotaxis, induction of CD44 expression on monocytes, and the adoption of a TAM gene signature in metastatic ovarian cancer.

Mice with EC-restricted *Rbpj loss* had impaired ovarian carcinoma growth in the omentum and peritoneum and lower numbers of MN-derived macrophages in the peritoneal fluid. These macrophages are essential for metastatic ovarian cancer development^4,30^. As such, the endothelium can influence tumour progression and metastasis by altering the immune status of the TME. Specifically, we observed that monocyte recruitment is potentiated, at least in part, through the release of *Rbpj*-mediated CXCL2 from ECs. Interestingly, higher serum levels of CXCL2 in ovarian cancer patients are associated with myeloid cell infiltration, poor prognosis and chemoresistance^31^. It should be noted that this chemokine has been traditionally associated with the recruitment of neutrophils^21^. However, there is evidence that CXCL2 also plays a role in the regulation of TAMs, especially those derived from monocytes. For instance, the CXCL2 receptor CXCR2 on monocytes and macrophages is important for the education of TAMs in prostate cancer^32^.

CXCR2 blockade has also been shown to re-sensitize ovarian cancer to cisplatin treatment^33^. Here we suggest that the CXCL2/CXCR2 axis might also have a role in the recruitment and education of macrophages in ovarian cancer. By separating newly recruited macrophages from macrophages that have reside in the peritoneal cavity for a longer period, we observed that endothelial RBPJ is necessary for the education into TAMs by tumour cells. Specifically, we report that MN-derived macrophages in contact with ECs lacking RBPJ had a lower expression of the HA receptor CD44. This receptor gets stimulated by HMW-HA produced by tumour cells to induce cholesterol depletion in macrophages, a crucial mechanism by which tumour cells educate TAMs^4^. Therefore, we propose that through this mechanism ECs can pre-condition MN-derived macrophages and contribute to their immunosuppressive phenotype within the tumour microenvironment. Indeed, the data showed that important genes involved in cholesterol depletion were reduced in MN-derived macrophages from mice lacking RBPJ in their ECs. Consistently, T cells isolated from this TME were more efficient at killing tumour cells, confirming the reduced immunosuppressive potential of the MN-derived macrophages.

In conclusion, we demonstrate that peritoneal ECs are critically involved in the recruitment and education of MN-derived macrophages in ovarian carcinoma.

## Material and methods

### Animal models

All animal procedures were approved by the local institutional animal care and use committee (RP Karlsruhe, Germany and DKFZ) and performed according to the guidelines of the local institution and the local government. Female C57BL/6 mice were group-housed under specific pathogen-free barrier conditions.

Administration of tamoxifen diluted in sterile peanut (P2144, Sigma-Aldrich, St. Louis, USA) in 8 to 12-week-old randomized mice was performed by oral gavage once with 100 μl (1 mg tamoxifen)^34^. Control mice, which did not express Cre^ERT2^ were also treated with tamoxifen.

Model of ovarian cancer: Three weeks after gene recombination, 5×10^6^ ID8-luciferase (ID8-luc) ovarian cancer cells were administered i.p. in PBS. For peritoneal lavage after sacrificing the mice, 5 ml of ice-cold PBS (Gibco/Thermo Fisher Scientific, NY, USA) was injected i.p., and after a careful massage to mobilize cells, peritoneal fluid was collected. For analysis of ID8-luc tumour growth the cell suspension was centrifuged and supernatant was collected. The cell pellet was suspended in 1 mL PBS and 100 μL were used to determine the luciferase activity. 100 μL cell suspension were centrifuged and the cell pellet was suspended in 100 μL lysis buffer (Promega) and 20 μL of lysed cells were pipetted into white 96-well plate in triplicates. 50 μL of LAR substrate (Promega) was added to the lysed cell suspension and luminescence signal was determined using the plate reader (ClarioStar, BMG Labtech). For analysis of immune cell recruitment into the peritoneal cavity the collected peritoneal cell suspension was centrifuged and red blood cells in the cell pellet were lysed with 1 mL ACK (Thermo Fisher Scientifics). After washing the cell suspension was counted using Neubauer Counting chamber and 1×10^6^ cells were used for flow cytometry staining.

Model of peritoneal inflammation: To obtain newly recruited peritoneal macrophages, 1 mL thioglycolate (2 mg/mL in H_2_O; B2551, Sigma-Aldrich) was injected into the peritoneum three weeks after gene recombination. MN-derived macrophages were isolated 24 hours after thioglycolate injection by their adherence to non-treated plastic petri dishes. Briefly, after incubation of single cells in a petri dish for 30 min at 37C, non-adherent cells washed away with PBS.

### Flow cytometry

For flow cytometry analysis, cells were suspended in 1 mL PBS with 2% FCS (Biochrom). Cell suspension was incubated with the different fluorophore-coupled primary antibodies for 20 min on ice. The following antibodies were used: CD45 (552848), CD11b (552850) and CD4 (560468) and CD8 (557654) from BD Biosciences (Bedford, MA, USA); CD3 (100203), F4/80 (123128), CCR2 (150608), CD44 (mouse and human 103007), CD74 (mouse (151005) and human (326811)) from BioLegend (St. Diego, CA, USA), MHCII (47-5321-80, Life Technologies/Thermo Fisher Scientific, NY, USA) and Tim4 (12-5866-82, Life Technologies/Thermo Fisher Scientific, NY, USA). Concentration of the different antibodies was determined by titration. In the meanwhile, compensation beads (UltraComp eBeads, Thermo Fisher) of the used primary antibodies were prepared. After staining, cells were washed with PBS and stored on ice until acquisition. Acquisition was performed using BD FACSCanto TM II, BD LSR for analysis or Aria for cell sorting (BD Biosciences).

Experiments were analysed using FlowJo Software. Flow cytometer results in percentage were extrapolated to the total number of cells obtained from the cell counting.

### Whole mount staining

The omentum was fixed for 1 hour in 1% PFA at room temperature and washed with PBS. Tissues were washed three times for 5 min with permeabilization buffer (PBS, 0.1% BSA and 0.2% Triton X-100) and treated with 0.5 mL blocking buffer (5% donkey serum diluted PBS-T) for 1 hour at room temperature. Primary antibodies, CD45 (BD 553076) and luciferase (ab185924), were diluted in PBS-T with 2% donkey serum. Next day, samples were washed three times with PBS-0.3% Tween. Afterwards secondary antibodies (1:200) were incubated for 1 hour at room temperature, washed three times with PBS 0.3% Tween and incubated with DAPI (1:10.000) for 15 min. Next, samples were incubated with clearing solution (FUnGi: 60% glycerol (vol/vol), 2.5M fructose, 2.5M urea, 10mM Tris Base, 1.0mM EDTA) at 4 °C overnight. Coverslips were mounted with FUnGi clearing solution and imaged with a confocal microscope (LSM 710, Carl Zeiss). All images were processed with ZENblue software (Carl Zeiss, Germany). Average mean intensities per image were counted with ImageJ software (NIH, Bethesda, MD, USA).

### Immunohistochemistry

Paraffin-embedded sections (3 μm) were de-paraffinized and re-hydrated in xylene and step-wise reductions in alcohol concentrations. H&E staining was performed according to standard protocols. Cytokeratin DAB and CD11b: antigen retrieval was performed at pH9 with citrate buffer. Primary antibodies CD11b (abcam, ab133357, 1:200), and Pan-cytokeratin (undiluted, ZUC001-125), diluted in blocking solution, were incubated at 4°C overnight. After washing, sections were incubated with secondary antibodies coupled with ZytoChem Plus (HRP) Polymer anti-mouse, (ZYTOMED, ZYT-ZUC050-006) and ZytoChem Plus (AP) Polymer anti-rabbit, (ZYTOMED, ZYT-ZUC031-006) diluted in antibody diluent (Cell Signaling) for one hour at room temperature. Afterwards, slides were treated with DAB Substrat Chromogen, Zytomed, ZYT-DAB057 and AP Red Kit, Zytomed, ZUC001-125. Immunofluorescence staining: primary antibodies: CD31 (Abcam, ab28364, 1:50), VCAM (Vcam, ab134047, 1:200) and αSMA (Sigma-Aldrich, A5228, 1:200) were diluted in blocking solution. Fluorophore-conjugated secondary antibodies (Thermo Fisher Scientific) were diluted in antibody diluent (Cell Signaling) together with isolectin-B4 (Thermo Fisher Scientific, I32450). Images were obtained with slide scanner (Zeiss Axio Sacn.Z1, Carl Zeiss) and a confocal microscope (LSM 710, Carl Zeiss). All images were processed with ZENblue software (Carl Zeiss, Germany). Image quantification were proceeded with ImageJ software (NIH, Bethesda, MD, USA).

### Cell culture

Murine cardiac ECs (MCEC) were purchased from tebu-bio and cultured on gelatin-coated surfaces in in DMEM containing 1 g/L D-glucose, 5 % FCS, 5 % HEPES, 100 units/ml penicillin and 100 μg/ml streptomycin.

HUVEC were grown and maintained until passage 5 in Endopan-3 Growth Medium containing 3% FCS and supplements (Pan-Biotech).

SK-OV-3 cells were kindly donated by Dr. Christiane Opitz (DKFZ, Heidelberg) and cultured in DMEM containing 1 g/L D-glucose, 10 % FCS, 5 % HEPES, 100 units/ml penicillin and 100 μg/ml streptomycin.

ID8-luc cells were kindly donated by provided by Prof. Frances Balkwill (Barts Cancer Institute, London, UK) and cultured in DMEM containing 1 g/L D-glucose, 10 % FCS, 5 % HEPES, 100 units/ml penicillin and 100 μg/ml streptomycin.

All cell culture experiments were performed in a laminar flow hood and cells cultured at 37°C and 95% relative humidity and 5% CO_2_. Cell culture were tested on a regularly basis for mycoplasma contamination periodically and before injecting the tumour cells into the mice.

To study the role of RBPJ in the regulation of Notch target genes, human umbilical vein endothelial cells (HUVECs) were infected with lentivirus constructs for CRISPR/Cas9 (Addgene plasmid #52961, lentiCRISPR v2) induced knock-out for *Rbpj^35^*. After 48 hours, cells were infected with adenovirus (LifeTechnologies; pAD/CMV-V5-DEST) overexpressing GFP as control or N1ICD.

### Isolation of peripheral blood nuclear cells (PBMC) from buffy coats

Human buffy coats were purchased from blood donation service DRK Mannheim, Germany. Peripheral blood nuclear cells (PBMC) were isolated by gradient centrifugation using Biocoll density solution (L6715; Biochrom). Human buffy coat was diluted 1:1 with PBS and added to the Biocoll density solution. This mixture was centrifuged at 430g for 20 min at room temperature. After centrifugation, the white intermediate phase containing leukocytes was collected and washed with PBS. To perform a positive isolation of monocytes, CD14 MACS beads were used with the LS column (130-042-402; Milentyi Biotec). The isolation of CD14^+^ monocytes was performed following the manufacture’s protocol.

### Transwell assay

Human ovarian cancer cells were seeded at 100,000 cells/ml in 500 μl RPMI medium without FCS for 48 hours. For the ECs monolayer, inserts were coated for 2 hours with 2 μg/ml fibronectin (1918-FN-02M; R&D Systems) in PBS. 50,000 human ECs *(RBPJ* knock-out or shRNA for *CXCL2* (pLKO.1; Sigma Aldrich) and respective controls) were seeded on top of the insert membrane for 48 hours. To analyse monocyte transmigration, 200,000 CD14^+^ cells were stained with carboxyfluorescein succinimidyl ester (CSFE; ThermoFischer) and added onto the endothelial monolayer. The transwell plate was incubated for 2 hours. For the chemotaxis assay with the recombinant proteins, transmigration of 50,000 CD14^+^cells were analysed towards 60 ng/mL CXCL2 (PeproTech GmbH) in RPMI medium without FCS for 30 min. After incubation, the remaining cell suspension in the upper well was aspirated and the transwell was cleaned with a cotton swab. The migrated cells were fixed with 4% PFA for 20 min at room temperature. Imaging of transwell was performed with Cell Observer (Carl Zeiss). From each transwell five evenly spaced field picture were taken using 20x objective and analysis was performed with Image J software.

### Enzyme-linked Immunosorbent Assay (ELISA)

Protein expression of CXCL2 (MIP-2) was quantified using an enzyme-linked immunosorbent assay (ELISA; Abcam ab184862). Cell culture supernatant was collected after 24 hours and ELISA was performed following the manufacture’s protocol.

### Bone marrow-derived macrophages (BMDMs) differentiation

Mouse macrophages were derived from the bone marrow of wild-type C57BL/6 mice. Femurs and tibiae were flushed several times with DMEM and collected cells were centrifuged. Bone marrow cells were suspended in media and seeded on 10 cm petri dishes (Corning). To differentiate these cells into macrophages, 10 ng/ml M-CSF (PeproTech GmbH) were added to Dulbecco’s modified Eagle’s medium (DMEM) (Thermo Fisher) supplemented with 10% fetal calf serum (FCS) (Biochrom, UK). Differentiation occurred within seven days. Cells were afterwards stimulated with indicated amount of recombinant protein CXCL2 (250-15-20, PeproTech GmbH) in DMEM medium without FCS or conditioned medium of endothelial cells.

### Immunostaining

BMDMs were cultured in DMEM with 10% FCS (Biochrom, UK) and 250,000 cells/well were seeded into 24-well plates on top of coverslips. BMDMs were treated with 40 ng/mL CXCL2 (PeproTech GmbH) in DMEM without FCS for 72 hours. Cells were washed with PBS and fixed with 4% PFA for 10 min. Then, the coverslips were washed three times for 5 minutes with PBS, permeabilized with PBS with 0.1 % Triton X-100 for 10 minutes and blocked for 30 min in blocking buffer (PBS in 5% FCS with 0.1% Tween 20 and 100 mM glycine) for 1 hour at room temperature. The coverslips were incubated with antibody against CD44 (1:1000) (Abcam, ab124515) and CD45 (1:500) (BD 553076) overnight at 4°C. The coverslips were rinsed three times with blocking buffer and incubated with a secondary antibody coupled to Alexa Fluor-488 and Alexa Fluor-546 (1:200) for 1 hour. The coverslips were washed and incubated with a DAPI solution before they were washed again. Coverslips were mounted and imaged with a confocal microscope (LSM 710, Carl Zeiss). All images were processed with ZENblack software (Carl Zeiss, Germany).

### cDNA synthesis and qPCR

RNA isolation from cell culture was performed using the InnuPrep Mini Kit (Analytik Jena) according manufacture’s protocol. RNA isolation form tissue was performed using PicoPure RNA Isolation Kit (Acturus, Life Technology). 1 mL Trizol was added to the tissue and homogenized for 1 min and a frequency of 30/sec. After disruption of the tissue, 200 μL chloroform were added and mixed by inverting several times followed by a centrifugation step for 15 minutes 12.000g at 4°C. Further RNA isolation steps were performed using the manufacture’s protocol. RNA concentration was measured using a Nanodrop 100 (Thermo Fisher Scientific). Reverse transcription of isolated RNA into complementary DNA was performed using High-Capacity cDNA Reverse Transcription Kit (Thermo Fisher Scientific).

Quantitative real-time PCR (qPCR) was performed with SYBR Green PCR mix (Applied Biosystems) on a QuantStudio3 Real-time PCR system (Applied Biosystems). Resulting fold changes were calculated using the 2^ΔΔCT^ method and mRNA expression was normalized to the housekeeping gene (*Cph* for murine and *HPRT* for human samples).

#### Primers used for mouse genes

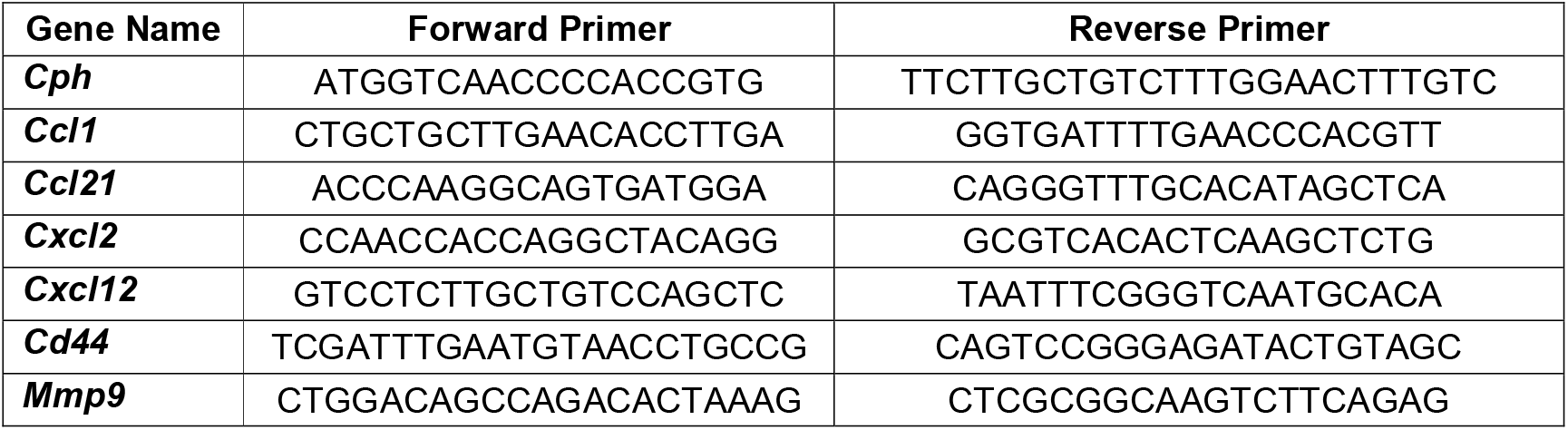

#### Primers used for human genes

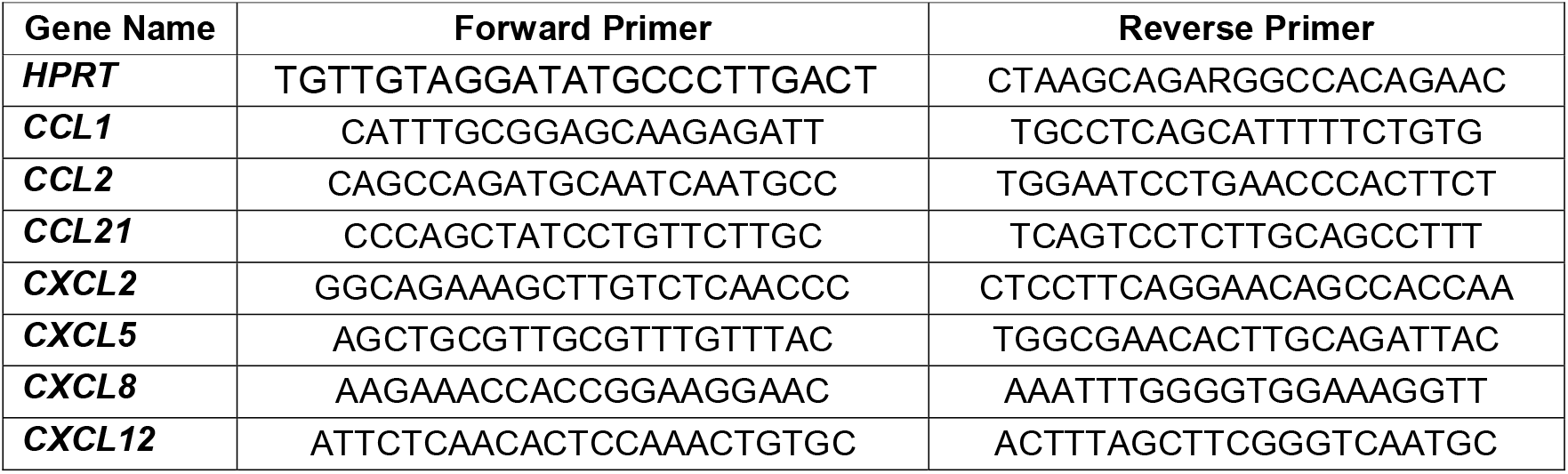

### Cytotoxicity assay

To analyse the T cell killing potential by the lactate dehydrogenase (LDH)-Cytotoxicity Assay Kit (Ab65393, Abcam), ID8 cells (7,500 ID8 cells in 100 μL DMEM medium) were seeded in a 96-well plate one day before T cell sorting. T cells were sorted from tumour bearing *Rbpj*^iΔEC^ and control mice after six weeks of tumour growth and 10,000 CD3^+^ cells were co-cultured with ID8 cells in technical triplicates including untreated control and blank. After overnight incubation, cytotoxicity and killing potential was measured by LDH amount in the cell supernatant using the LDH-Cytotoxicity Assay Kit (Ab65393, Abcam) following the manufacturer’s protocol.

#### *In silco* analysis of promotor region

To determine RBPJ binding sites^36^ (5’-GTGGGAA-3’) in the promotor region of the murine (NM_009140; chr5+:90902580-90903927) and human (NM_002089; chr4-:74100502-74099123) CXCL2-encoding gene, we used ApE plasmid Editor by M. Wayne Davis (https://jorgensen.biology.utah.edu/wayned/ape/).

#### *In silico* protein-protein interaction

We used the Search Tool of Interacting Genes/Proteins database (STRING v11.5) to perform *in silico* protein-protein interaction analysis^37^. Given CXCR2 protein as input, STRING can search for their neighbour interactors, the proteins that have direct interactions with the inputted proteins; then STRING can generate the PPI network consisting of all these proteins and all the interactions between them. All the interactions between them were derived from high-throughput lab experiments and previous knowledge in curated databases at medium level of confidence (score ≥ 0.40).

### Gene set enrichment analysis (GSEA)

GSEA (Broad Institute) was used to determine if a list of genes (gene signature) was enriched between different groups. A defined list of genes exhibits a statistically significant bias in their distribution (false discovery rate (FDR)) within a ranked gene list using the software GSEA^38^ resulting to an enrichment in one of the compared groups (normalized enrichment score (NES)) obtained from microarray or public available data sets as indicated in the figure legend.

### Ingenuity Pathway Analysis (IPA)

IPA software (Qiagen) was used to identify predicated upstream regulators and differentially regulated pathways in newly recruited macrophages (based on microarray data). For the analysis of data, fold-changes were uploaded to the software. Differentially regulated pathways and upstream regulator analysis was performed from obtained microarray data.

### Pathway analysis

Pathway analysis were obtained from public external databases (EnrichR) and analysed as −2log fold changes.

### Human ovarian cancer patient RNAseq data analysis

Human ovarian cancer patient bulk tumour RNA-sequencing data was obtained from the Cancer Genome Atlas (TCGA) database. Stratification in CXCL2^high^ and CXCL2^low^ patients was performed using R Studio software. Patients were assigned to the different groups using *CXCL2* expression below the first or higher than the third quartile. Normalised raw counts of CD44 were plotted comparing the two different groups.

### Statistical analysis

Normality was tested when sample size allowed it. Those samples with normal distribution where compared using Students’ t-test (with Welch’s correction when groups had different sizes). When normality was rejected, Mann-Whitney U-test was used. Comparison analysis was performed with analysis of variance (ANOVA) with Tukey post-test when more than two groups were analysed. Statistical analysis and the generation of the graphs were performed using GraphPad Prism 9 (GraphPad Software, Inc.; La Jolla, CA, USA).

### Schematic Figures

Schematics were created using BioRender.com.

## Acknowledgements

We thank Ralf Adams (MPI Münster, Germany) for providing Cdh5-CreERT2 mice, we kindly acknowledge Dr. Christiane Opitz for providing the SK-OV-3 cell line and Frances Balkwill (Barts Cancer Institute, London, UK) for providing ID8-luc cells. We thank the Light Microscopy core facility, the Microarray Unit, the Flow Cytometry core facility and animal caretakers of the German Cancer Research Center (DKFZ) for providing excellent services. We would like to thank Damir Krunic (DKFZ, Light Microscopy Core Facility) in particular for his help with FIJI software data analysis.

This work was funded by the Deutsche Forschungsgemeinschaft (DFG) project 394046768 - SFB1366 projects C4, C2 (to A.F.& A.C.), SPP 1937 (CE 140/2-2 to A.C.), TRR179 (TP07 to A.C.), SFB-TRR156 (B10N to A.C.); the Cooperation Program in Cancer Research of the German Research Cancer Center (DKFZ) and the Israeli Ministry of Science and Technology (MOST) (to A.F.), DFG project 419966437, Deutsche Krebshilfe project 70113888, MCIN/AEI/ 10.13039/501100011033 (PID2020-117946GB-I00 and RYC2019-027937-I) (to J.R-V.) and by “ERDF A way of making Europe” program and by “ESF Investing in your future”. Part of the equipment used in this work has been funded by Generalitat Valenciana and co-financed with ERDF funds (OP ERDF of Comunitat Valenciana 2014-2020).

## Author Contributions

R.M., E.A-S, I.M., S.B., L.W., L.J., T.Z., J.T., F.A.R., A.S., B.D.G., J.R-V. performed experiments. A.C., T.B., A.F. and J.R-V. contributed to analysis of results. A.F. and J.R-V. conceived the original hypothesis. R.M., E.A-S., A.F. and J.R-V. planned experiments. R.M., E.A-S., J.R-V. and A.F. wrote the manuscript. J.R-V. directed and supervised the work.

