## Supplemental data for "Endothelial Rbpj is essential for the education of tumour-associated macrophages"

**Supplement Figure 1. Effects of loss of endothelial *Rbpj* in omental endothelium**

**a,** Representative microscopic images of omentum stained with H&E from non-tumour bearing *Rbpj*$^{i\Delta EC}$ and control mice. Scale bar, 1mm. **b**, Representative images of immunohistochemistry staining for CD31 (white) and DAPI (blue) in omentum of non-tumour bearing *Rbpj*$^{i\Delta EC}$ and control mice. Scale bar, 50 µm. Quantification of vessel density (relative CD31+ area). Bar graphs show mean±SD; two-tailed, unpaired Mann-Whitney U-test **c,** Representative images of immunohistochemistry staining for CD31 (white), αSMA (red) and DAPI (blue) in omentum of *Rbpj*$^{i\Delta EC}$ and control mice after four weeks of ID8 tumour growth. Scale bar, 100 µm. Quantification of vessel coverage (CD31+SMA+ area). Bar graphs show mean±SD; two-tailed, unpaired Mann-Whitney U-test.

**Supplement Figure 2. Effects of loss of endothelial *Rbpj* on myeloid cell infiltration under physiological conditions**

**a,** Representative images of immunohistochemistry staining for CD31 (white), VCAM1 (red) and DAPI (blue) in omentum of non-tumour bearing (basal) and four weeks after tumor growth from *Rbpj*$^{i\Delta EC}$ and control mice. Scale bar, 50 µm. Quantification of n≥3. Bar graphs show mean±SD; two-tailed, unpaired Mann-Whitney U-test. **b**, Full representative images of whole mount staining of tumour nodules of the omentum for luciferase (red), CD45 (green) and DAPI (blue) four weeks after tumour injection in *Rbpj*$^{i\Delta EC}$ and control mice. Scale bar, 20 µm. **c**, Myeloid cells ($CD45^{+}$, $CD11b^{high}$) and **d,** Macrophages ($CD45^{+}$, $CD11b^{high}$, $F4/80^{+}$) relative to alive cells. n=3. Bar graphs show mean ±SD; two-tailed, unpaired Mann-Whitney U-test. **e** and **f,** macrophage subpopulations characterization by F4/80 and MHCII expression into LPM and SPM in *Rbpj*$^{i\Delta EC}$ mice compared to controls and their quantification. n=3. Bar graphs show mean ±SD; two-tailed, unpaired Mann-Whitney U-test.

**Supplement Figure 3. RBPJ-independent Notch1 regulation of cytokines in endothelial cells**

Quantification of mRNA expression of *CXCL5, CXCL8* and *CCL2* upon NICD overexpression, knock out of *RBPJ* and combination in HUVEC. n≥3. Bar graphs show n-fold vs GFP transduced as mean±SD; two-tailed, unpaired Mann-Whitney U-test.

**Supplement Figure 4. Essential role of endothelial *Rbpj* in regulating CXCL2 mediated expression of CD44 on macrophages**

Full representative images of BMDMs stained with CD44 (red), CD45 (green) and DAPI (blue) of control and stimulated with CXCL2 (40 ng/mL).

**Supplement Figure 5. Gene ontology (GO)-term analysis in TCGA patients**

Gene ontology (GO)-term analysis, biological processes (left) and molecular function (right) of CXCL2high patients using R Studio software (R Core Team (2017)); n=97.

**Supplement Figure 6. No changes in CD74-mediated gene signature for resident peritoneal macrophages (CCR2$^{-}$).**

GSEA of resident macrophages (CD45$^{+}$; CD11b$^{+}$; F4/80$^{+}$; CCR2$^{-}$) from *Rbpj*$^{i\Delta EC}$ and control mice compared with CD74-mediated gene signature with 10 most differentially regulated gene extracted from TAM signature of ID8 tumor growth.

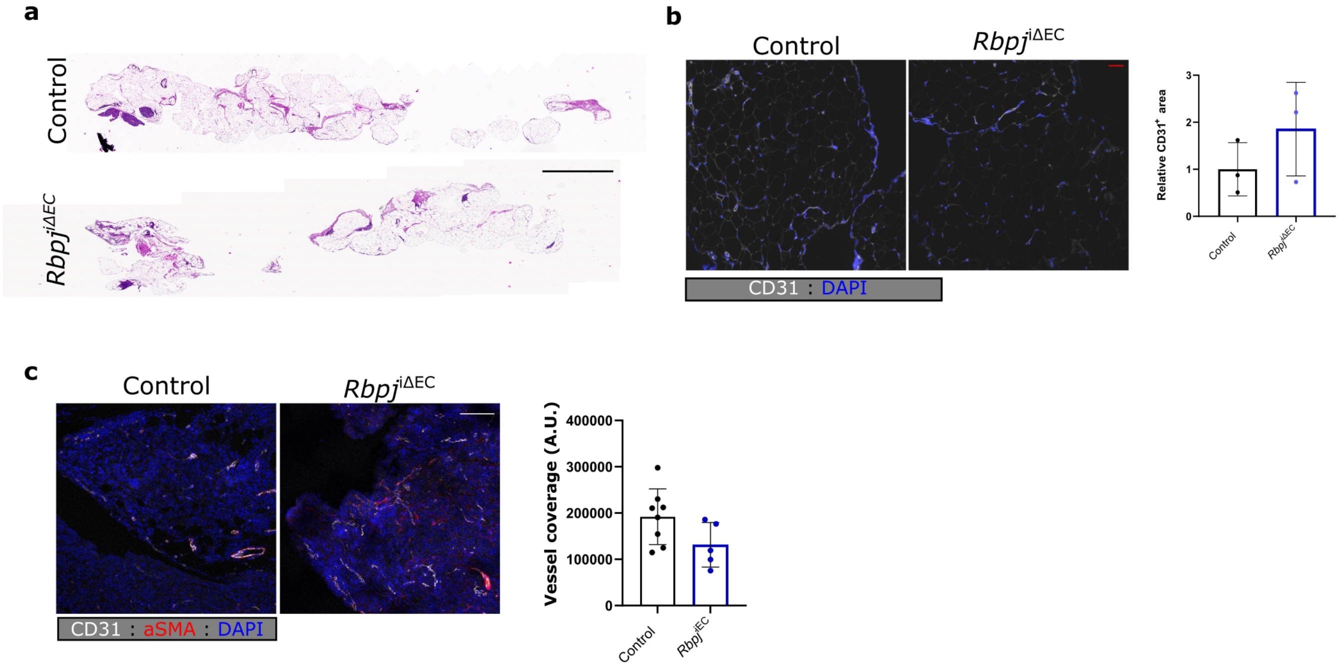


**Supplementary figure 1.**

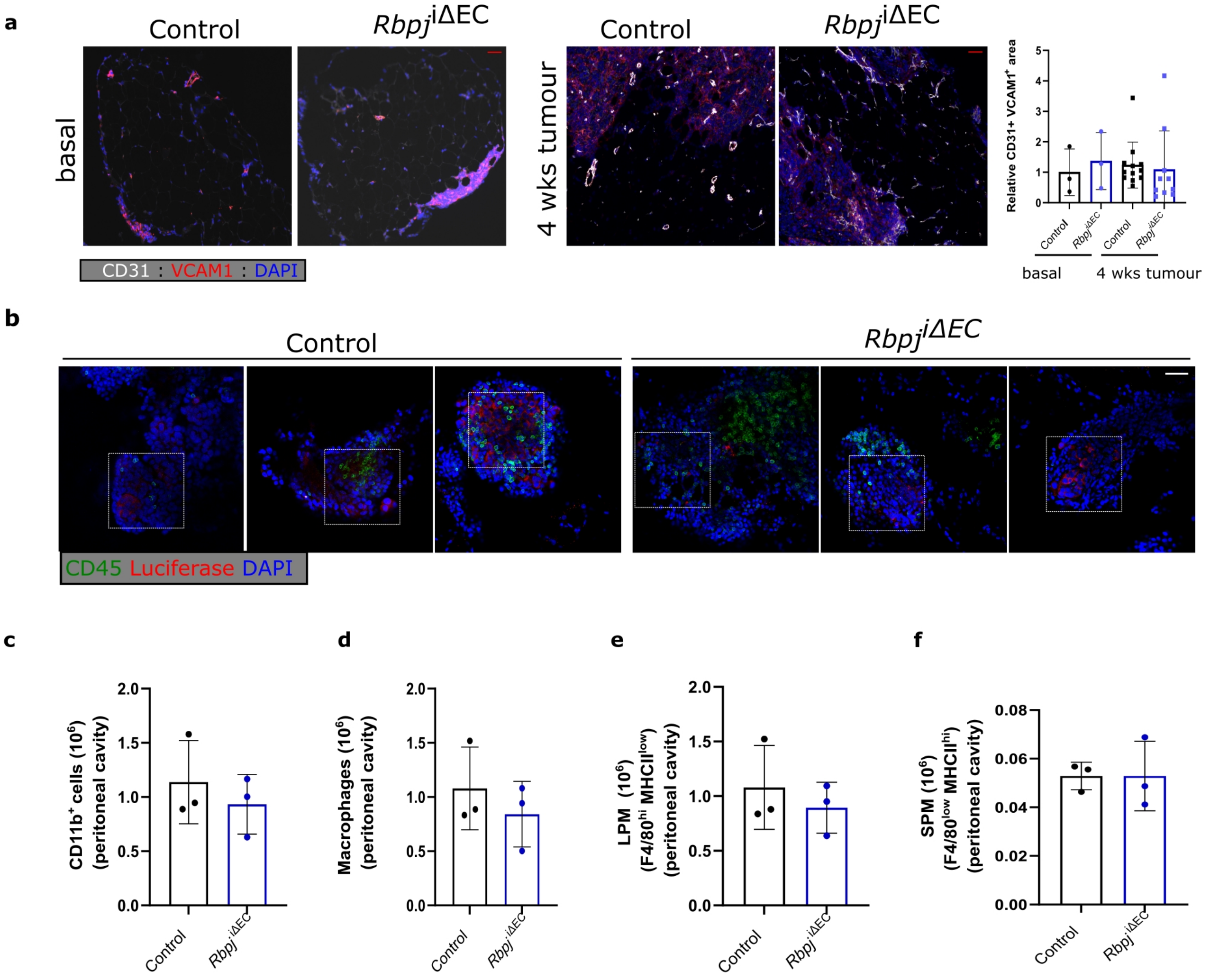


**Supplementary figure 2**.

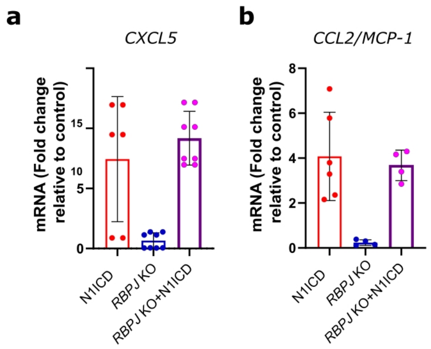


**Supplementary figure 3**.

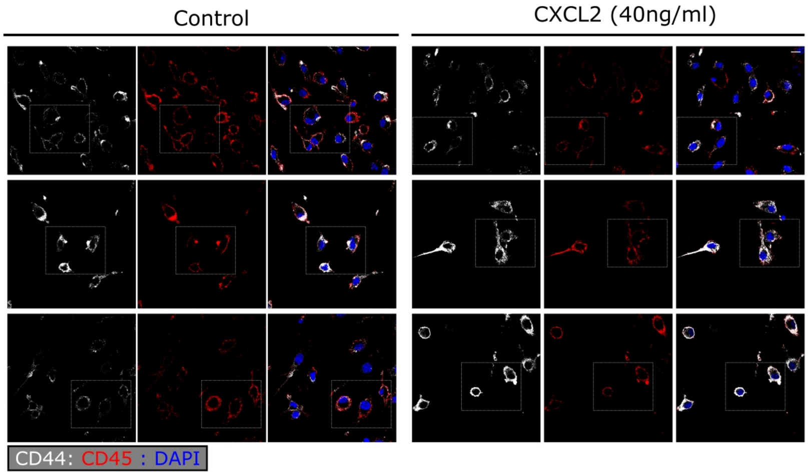


**Supplementary figure 4.**

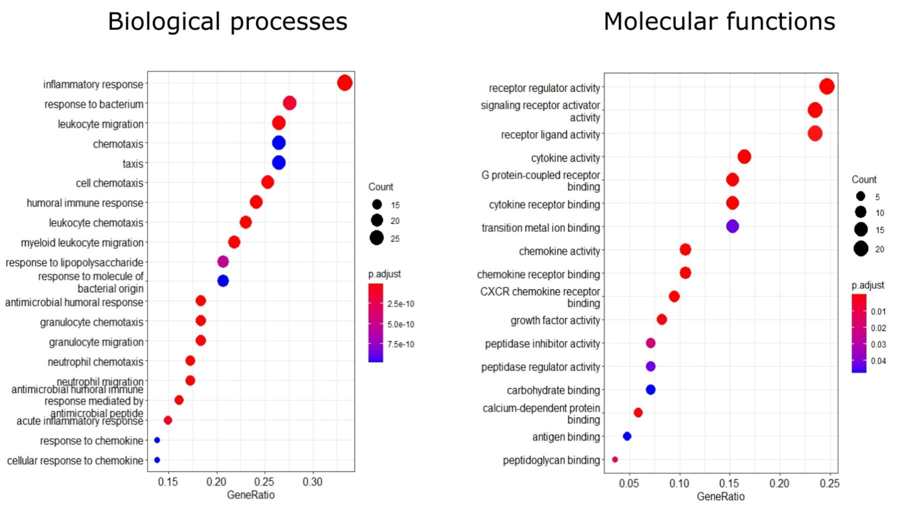


**Supplement Figure 5**.

Resident Macrophages

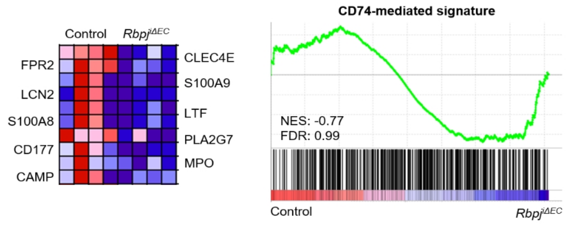


**Supplementary figure 6.**
